# Facial expression discrimination emerges from neural subspaces shared with detection and identity

**DOI:** 10.1101/2025.08.25.672186

**Authors:** Maren Wehrheim, Shirin Taghian Alamooti, Hamidreza Ramezanpour, Kohitij Kar

## Abstract

Understanding how the human brain decodes facial expressions remains a fundamental challenge, requiring computational models that tightly connect neural responses to behavior. Here, we demonstrate that rhesus macaques provide a unique and powerful animal model to uncover the neural computations behind human facial expression discrimination, bridging critical gaps between behavior, neural activity, and computational theory. Despite the challenges of establishing reliable behavioral paradigms in macaques, we developed a robust discrimination task spanning six emotional categories, yielding strong, image-by-image behavioral correspondence between macaques and humans. By systematically comparing artificial neural networks (ANNs) to macaque behavior and IT neural data, we found that traditional action unit–based models fail to capture image-level behavioral structure, while ANNs with IT-like internal representations outperform all others. Neural recordings showed that the specific IT population responses (70–100 ms) carried the strongest predictive power for facial expression discrimination, underscoring the primacy of feedforward codes in guiding behavior. Expression coding in IT was significantly shaped by face-selective neurons that also encoded identity. This convergence points to a shared functional subspace in IT, where stable (identity) and dynamic (expression) information coexist along overlapping dimensions. Such an architecture moves beyond the classical view of segregated pathways, revealing a general coding principle by which IT flexibly supports multiple socially relevant functions within a common representational geometry.

## Introduction

Discriminating facial expressions is fundamental to human social behavior, yet our understanding of the neural mechanisms driving this perceptual ability remains incomplete^1,2^. A significant barrier has been the lack of rigorous animal models that simultaneously capture precise human behavioral outcomes and the corresponding neural mechanisms at a resolution necessary to guide biologically realistic computational models. Rhesus macaques represent a particularly compelling model, given their extensive anatomical and functional parallels to human visual processing systems, notably within the inferior temporal (IT) cortex, critical for object^3^ and face recognition^4,5^.

Previous research in both humans and macaques highlights overlapping neural substrates within the temporal cortex specialized for processing facial identity and emotional expressions^6,7^. Neurons in macaque IT cortex selectively encode facial features essential for recognizing identity^8^ and emotional states^9–11^, and behavioral data^12^ confirm that macaques respond to facial emotional cues analogously to humans. Despite these parallels, existing macaque studies have largely relied on qualitative or simplistic behavioral paradigms and have predominantly used conspecific (macaque) faces^13^. This methodological choice significantly limits direct inferences regarding human neural mechanisms. Consequently, a critical unresolved issue is whether macaques can serve as accurate behavioral and neural models for the purely sensory aspects of human facial expression discrimination, specifically at the level of individual human facial images. Without rigorous validation of this behavioral alignment, the potential of macaques to inform computational hypotheses about human facial emotion recognition remains uncertain.

Moreover, even if detailed neural-behavioral patterns are observed in macaques, an essential step remains: how can such information be leveraged to update our understanding of human neural mechanisms? Ultimately, a key goal needs to be to translate insights from animal models into computational hypotheses testable in humans. Specific artificial neural networks (ANNs), which have emerged as highly predictive computational models of the primate ventral visual pathway, provide a powerful tool for this translation^3,14^. ANN models of the ventral visual stream have shown strong alignment with primate neural responses^15–17^ in object recognition and can partially account for neural coding in face-selective regions for identity processing^18^. Similarly, action unit–based approaches provide a structured, interpretable framework for facial expression recognition^19,20^. However, recent work indicates that both approaches leave substantial explainable variance unexplained, particularly for socially meaningful, fine-grained judgments such as facial expressions^21^. This suggests that while these models capture some representational structure, their ability to account for the neural computations underlying facial expression discrimination remains largely untested. Thus, insights derived from macaque behavioral and neural data have the potential to critically guide the improvement of these computational models, refining their biological alignment and, consequently, their utility in modeling human facial expression discrimination.

To address these interconnected gaps explicitly, our study poses three core questions: (1) Can macaques reliably discriminate between human facial expressions, and to what extent do their image-specific behavioral patterns align with those of humans? (2) What computational hypotheses about neural processing do existing ANN models provide and can these models explain the observed macaque behavioral patterns? (3) Can neural responses from the macaque IT cortex directly predict macaque behavioral discrimination patterns, and if so, what roles do neuronal selectivity, identity–expression representational overlap, and specific temporal windows of processing play in driving this predictive power?

We first established macaques as a valid model organism for studying the mechanisms of human facial expression discrimination using a comprehensive set of human facial expression stimuli^22^ (see examples in Fig. 1A) and rigorously quantifying the behavioral alignment at the image level between macaques and humans (Fig. 1B). Specifically, we developed a robust behavioral paradigm (Fig. 1C) and associated behavioral metrics (Fig. 1D) in macaques, explicitly designed for direct and quantitative comparison with human performance. Subsequently, we systematically evaluated multiple ANN architectures, including models specialized for facial action units, and ANNs with varied architectures, objectives, and learning strategies that best predict primate neural data, by comparing their facial expression discrimination behavior and the internal representations against primate behavior and neural responses recorded from the macaque IT cortex respectively (Fig. 1E). Critically, this allowed us to determine which computational approaches best explain the neural encoding strategies underlying facial expression discrimination. Guided by hypotheses from ANN analyses, we next linked direct neural recordings from macaque IT cortex to detailed image-level behavioral performance to test whether population activity could robustly predict expression discrimination behavior. Finally, we assessed whether neurons’ face selectivity and their encoding of facial identity accounted for their contribution to expression discrimination, revealing how these properties shape behavioral alignment at both the population and single-neuron levels.

**Fig. 1:**
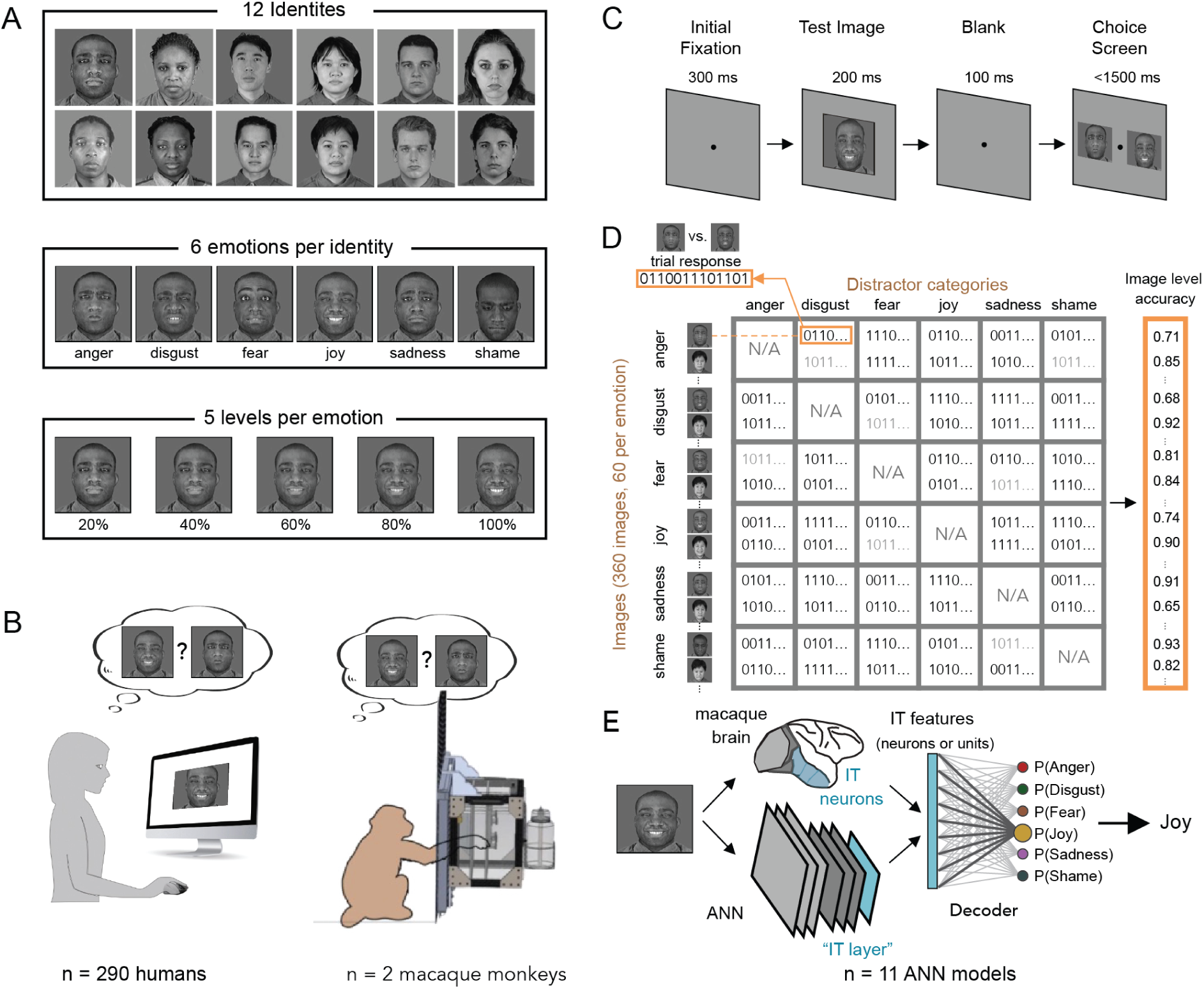
Experimental and computational framework for studying facial expression discrimination. A) Stimulus Set. The imageset comprises twelve identities (balanced for gender and ethnicity) each displayed six facial emotions at five graded intensities, yielding a stimulus set designed to probe both categorical and image-level variability. **B) Behavioral testing across primates.** Humans (left) perform the facial expression discrimination task online via Amazon Mechanical Turk, while macaque monkeys (right) were tested on the same task using custom-built kiosks^25^ that were attached to their home cages. **C) Task paradigm.** On each trial, a *Test* image of a face with a specific emotional expression (at a specific graded intensity) is presented for 200 ms, followed by a 100 ms delay. Subjects then choose the expression of the *Test* image (the *Target* expression category) performing a forced choice between the correct expression and a distractor expression, both shown at 100% intensity for the same identity. **D) Image-level accuracy computation and the B.I1 performance metric**. Binary trial responses (1 for correct, and 0 for incorrect) for each Test image and distractor expression are averaged to first create an image x distractor matrix. Averaging across all distractors generates an accuracy score for each image individually (the B.I1 vector), enabling quantitative comparisons across humans, macaques, and models. **E) Neural and ANN prediction.** For each image, we extracted responses from macaque IT cortex and from IT-like layers of artificial neural networks (ANNs). For each, a linear one-vs.-all classifier (decoder) is trained to predict emotion labels, and the resulting predictions were compared against observed behavioral performance.

Importantly, our study specifically targets the perceptual and sensory aspects of facial expression discrimination, isolating them from emotional or affective processing. By using brief stimulus presentations followed by immediate forced-choice discrimination of the expression within the same facial identity, we partially restrict that performance reflects visual encoding rather than higher-order cognitive or affective evaluation^23^. This targeted approach enables a rigorous examination of the visual neural mechanisms underlying the macaque’s processing of human facial expression, likely free from emotional or experiential influences, unlike studies that use conspecific faces^24^.

Building on this foundation, our integrative approach—combining precise human and macaque behavioral paradigms, large-scale recordings from macaque IT cortex, and ANN modeling—directly links neural population codes to behavior and model predictions. This framework allowed us to test whether macaques replicate human image-level performance, to evaluate which ANN architectures generate biologically meaningful hypotheses, and to determine which features of IT activity predict behavioral outcomes. Crucially, our results reveal that expression discrimination depends most strongly on early-phase IT population responses and on neurons that are both face-selective and identity-informative, pointing to a shared representational subspace where stable and dynamic facial features coexist. This organization moves beyond the canonical view of segregated face pathways, instead defining a general coding principle by which IT flexibly supports multiple socially relevant functions. More broadly, the combination of cross-species behavior, ANN modeling, and neural population recordings provides a generalizable blueprint for dissecting how high-level visual cortex transforms sensory inputs into behaviorally relevant codes.

## Results

As outlined above, we begin by developing a human facial expression discrimination task that can be tested without significant modifications across humans, macaques, and ANNs. We designed our behavioral paradigm with several key considerations in mind to ensure that it would capture sensory-level neural computations and remain compatible with both macaque and human testing. First, we included a wide range of emotional expressions—six distinct categories (anger, disgust, fear, joy, sadness, shame)—across 12 identities of varying gender and ethnicity. This diversity allowed us to assess performance across a broad stimulus space and to examine image-specific effects. Second, we kept stimulus presentation brief (200 ms), followed by a short delay (100 ms) before the response phase to minimize the influence of higher-order cognitive strategies, keeping the task closely tied to purely perceptual processing. Third, we used a relatively large number of images per category, enabling image-level behavioral metrics that could be directly compared with image-computable ANN outputs. Fourth, we controlled low-level image properties (luminance, contrast, and spatial frequency) using the SHINE toolbox to ensure that performance differences could not be attributed to trivial low-level cues. Together, these design choices allowed us to create a task that was both biologically relevant and computationally tractable, serving as a rigorous benchmark for human–macaque comparisons and ANN evaluation.

### Consistent Human Performance with Systematic Variability Across Emotions

We first sought to establish whether human participants exhibited the behavioral properties necessary for this paradigm to serve as a rigorous benchmark for cross-species comparisons. Specifically, we asked whether image-level accuracy varied sufficiently across stimuli within each emotion category, and whether these patterns were reproducible across participants—both of which are critical for assessing whether macaques (and later, ANN models) can capture the fine-grained structure of human performance. A total of 290 human subjects on Amazon Mechanical Turk (see Methods for quality control) completed the expression discrimination task under identical timing constraints to those planned for the macaque experiments (200 ms Test stimulus, 100 ms delay, two-alternative forced choice; see Fig. 1C).

As a first validation step, we computed the psychometric functions for each emotion category. As expected, we observed that discrimination accuracy increased monotonically with emotion intensity, reflecting the greater availability of visual cues at higher intensities—a simple but important sanity check. All emotion categories showed this trend (all slopes > 0), confirming that the task reliably captured graded sensitivity to expression strength (Fig. 2A).

**Fig. 2:**
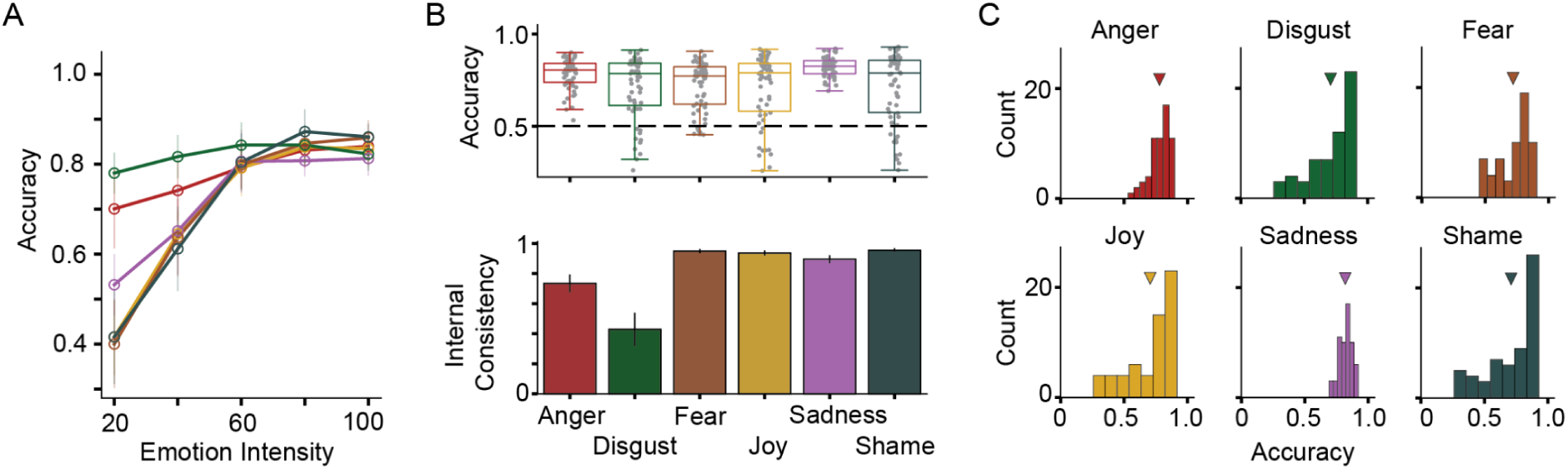
Human behavioral benchmarks for facial expression discrimination. **A) Psychometric validation.** Psychometric functions for each expression category (color-coded), showing mean accuracy as a function of expression intensity. Accuracy increased monotonically with expression intensity for each category, confirming that the task elicited expected psychometric behavior. Error bars indicate ±1 standard deviation across images. **B) High accuracy and consistent B.I1 patterns across emotions.** Image-level accuracy (top) and internal consistency (bottom) for each emotion. Each dot indicates an image and the box shows the mean and percentiles across images. The error bar for internal consistency shows the 95% confidence interval. **C) Distribution of image-level accuracies across each emotion category.** Distributions of image-level accuracy revealed similarly skewed patterns across all emotions, indicating that variability in discriminability is an intrinsic property of the stimulus set rather than category-specific. The triangle indicates the average performance per emotion.

We next examined whether accuracy varied across images within each category. Image-level accuracy scores spanned a wide range (0.7 – 0.82; where chance-level is 0.5), indicating that some expressions were consistently easier to discriminate than others, even at matched intensity levels (Fig. 2B–C). This variability is essential for comparisons across species at the image level. To ensure these patterns reflected robust perceptual signatures rather than noise, we computed internal consistency measures across participants. High consistency values were observed for all emotions (*r_SB_* ≥ 0.73), with the exception of a modest drop for “Disgust” (*r_SB_* = 0.43), confirming that the observed performance patterns were reproducible. Having established that human behavior in this task displays the required reliable variance across images, we next asked whether macaques tested on the same stimuli and paradigm show comparable performance patterns.

### Macaques Achieve High, Reliable Performance in Human Facial Expression Discrimination

Training macaques to discriminate emotional expressions in human faces required multiple stages (see Methods), as this task demands generalization across unfamiliar identities and detection of image changes generated by subtle facial muscle movements. Following training, both monkeys performed the task well above chance levels (*M* = 0.65, Wilcoxon signed-rank test *W* = 60217.0, *n* = 360, *p* < .001). Consistent with the human behavioral results, their performance showed: (a) human-like psychometric functions (Fig. 3A), with accuracy increasing with emotion intensity for most categories (all slopes > 0); (b) varied image-level accuracy patterns (Fig. 3B; top panel), reflecting differences in relative difficulty across stimuli; and (c) high internal consistency (*r_SB_* = 0.75 - 0.98) across emotions), confirming that these patterns were stable across repeated presentations (Fig. 3B; bottom panel). Furthermore, the two monkeys exhibited highly similar image-level accuracy patterns (noise corrected Pearson ρ = 0.97, *p* < 0.001, Fig. 3C), indicating consistent strategies within the monkeys. These results establish that macaques can accurately perform the human facial expression discrimination task and produce structured, reproducible behavioral patterns.

**Fig. 3:**
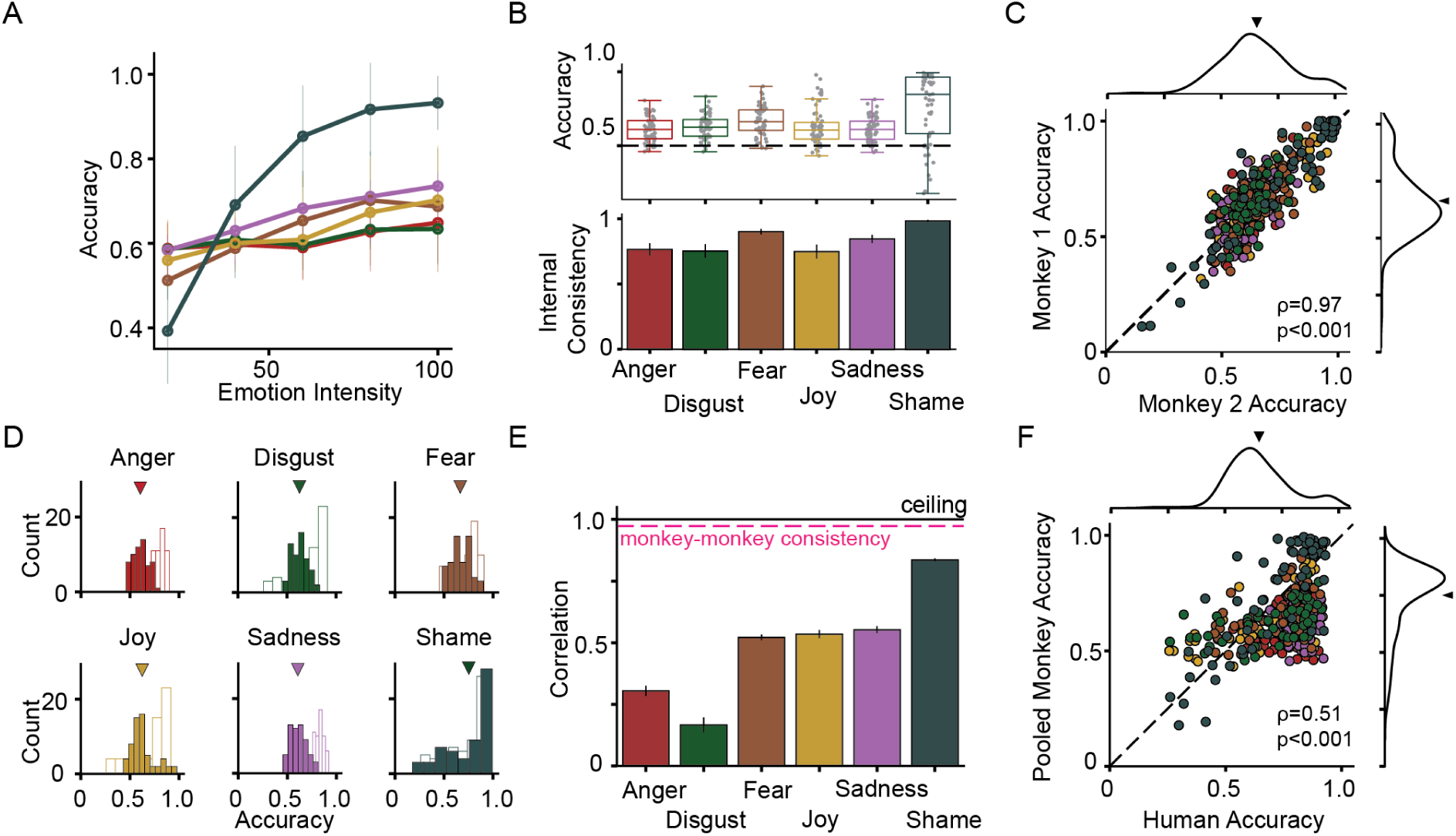
Validating macaques as a behavioral model for human facial expression discrimination. **A) Psychometric validation.** Psychometric functions for each emotion category (color-coded), showing mean accuracy as a function of emotion intensity. Accuracy increased monotonically with expression intensity for each category, confirming that the task elicited expected psychometric behavior. Error bars indicate ± 1 standard deviation across images. **B) Robust and reliable performance across emotions.** Image-level accuracy (top) and internal consistency (bottom) for each emotion. Macaque performance was above chance for all emotions. Each dot indicates an image and the box shows the mean and percentiles across images. The error bar for internal consistency shows the 95% confidence interval. **C) Cross-animal consistency.** Image-level accuracy for each monkey, showing a strong correlation across stimuli (ρ = 0.97, p < 0.001), indicating that behavioral patterns were reproducible across individuals, and that the training paradigm resulted in similarly trained macaques. Each dot represents a single image, colored by emotion category. The density plots show the monkey-specific distribution across all images and the triangle indicates the mean accuracy. The dashed line represents the bisector (x=y line). **D) Distribution of image-level accuracies.** Macaque (solid fill) and human (no fill) distributions of image-level accuracy for each emotion. The triangle indicates the macaque average performance per emotion. We observed that mean performance is significantly lower in macaques compared to humans across emotion categories (except for shame). **E) High emotion-specific performance overlap between species.** Each bar represents the trial repetition noise corrected correlation between the emotion-specific image-level performance patterns between the pooled monkey (n=2) and pooled human data. The error bar indicates the 95% confidence interval across different bootstrapped samples. **F) High cross-species image-level correlation.** Pooled macaque overall image-level accuracy patterns correlated strongly with human accuracy (ρ = 0.51, p < 0.001). Each dot represents a single image, colored by emotion category. The density plots show the monkey-specific distribution across all images and the triangle indicates the mean accuracy. The dashed line represents the bisector (x=y line).

### Macaque and Human Performance Patterns Are Strongly Correlated Despite Accuracy Differences

We next compared macaque performance directly to that of human participants to quantify cross-species similarity. While overall macaque accuracy was significantly lower than that of humans (Fig. 3D, Wilcoxon signed-rank test: *W* = 12549, *n* = 360, *p* < .001), such a difference was expected given species-specific familiarity with human faces. Distributional comparisons revealed that macaque performance was less skewed toward high accuracies than human performance, except for “Shame,” where the two species overlapped closely. The more critical question was whether both species shared the same relative difficulty structure across stimuli. Indeed, expression-specific correlations (corrected for the internal consistencies of humans and macaques) were high for most categories (range of noise corrected Pearson ρ = 0.18–0.84; Fig. 3E). These correlations were significantly above chance (*p* < 0.001 permutation test) and comparable to cross-species similarity levels reported for object recognition^26,27^. At the overall image-level, the pooled macaque accuracy patterns correlated strongly with human image-level accuracies (noise corrected Pearson ρ = 0.51, *p* < 0.001; Fig. 3F). Together, these findings indicate that while macaques perform at a lower absolute accuracy, their behavioral patterns across images and emotional expressions closely parallel those of humans, validating them as a behavioral model for human facial expression discrimination.

### ANNs Reveal Gaps in Behavioral Alignment and Opposing Temporal Predictions

We next turned to artificial neural network (ANN) models of the primate ventral stream to generate computational hypotheses about the mechanisms underlying the observed macaque facial expression discrimination behavior. As with the behavioral analyses above, we began by asking whether these models could perform the task (Fig. 4A). All tested ANN models achieved near-perfect classification accuracy on our stimulus set, with many substantially overperforming relative to macaques (*M* = 0.88, range: 0.64–0.93; Fig. S1A). To enable fair comparisons of behavioral patterns, we therefore randomly subsampled features from each model until its accuracy matched the pooled macaque level (see Methods, Fig. S1B).

**Fig. 4:**
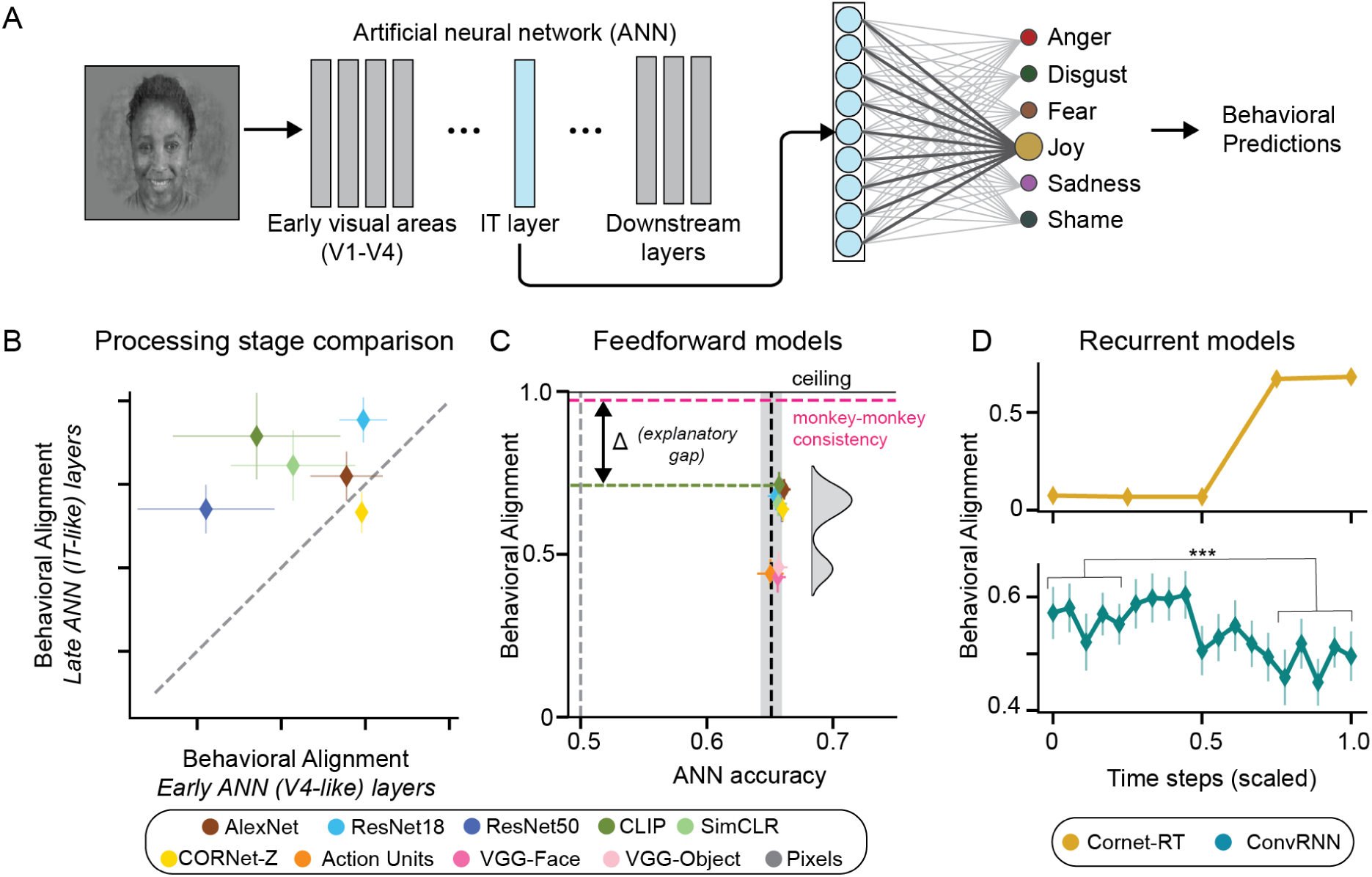
Persistent gap between primate behavior and artificial neural networks. A) Prediction pipeline. For each ANN, we extracted activations from a given layer (Brain-Score labelled IT layer shown in blue) and used it to train six, cross validated one-vs-rest logistic regression classifiers to predict expression category labels. The ANN units from these layers were subsampled until classification accuracy matched pooled macaque accuracy, ensuring fair behavioral alignment comparisons across ANNs. Model predictions were then compared to macaque image-level behavioral accuracy patterns. **B) Late layers better predict behavior.** Later, IT-like layers (deepest 30% of layers) of static models showed stronger alignment with macaque image-level behavior than earlier layers (first 30% of layers), consistent with hierarchical ventral stream processing. Each dot represents the mean across layers within the early or late group. Error bars show the standard deviation across layers. The dashed line marks the bisector. **C) Behavioral explanatory gap despite matched accuracy**. Using only the IT-aligned layer for each ANN, each dot shows the mean results across 50 iterations; horizontal error bars show the standard error of mean accuracy across images, vertical error bars show the standard deviation of 100 bootstrap images. The vertical dashed line and shaded area indicate the mean and standard error of macaque accuracy. The magenta dashed line shows the consistency of one monkey’s image-level accuracy with the other. The distribution of ANN-behavior alignment scores is shown on the right. **D) Divergent recurrent dynamics across convRNN and CORnet-RT.** Corrected behavioral alignment is shown for each time frame, with time normalized between 0 and 1 for comparability across models. CORNet-RT showed increasing behavioral alignment over time, whereas ConvRNN alignment declined significantly in later time bins, despite both achieving similar accuracy levels. These contrasting profiles provide distinct hypotheses for neural dynamics. Error bars show the standard deviation of 100 bootstrap images.

After matching the average model accuracy across images to the average macaque levels through feature subsampling, we examined how behavioral alignment varied across the ANN space. Consistent with the idea that macaque facial expression discrimination relies on higher-order ventral stream computations, we found that features from later, IT-like layers aligned more closely with macaque behavioral patterns than features from earlier layers (*t*(4) = -2.73, *p* < 0.05, *d* = -1.57; Fig. 4B). However, we observed that even the best-performing layers failed to fully reproduce macaque image-level behavior. Despite identical overall accuracies, alignment at the image-level varied widely across models (max corrected correlation = 0.71), revealing an explanatory gap between model representations and the behavioral structure observed in macaques (Fig. 4C). Interestingly, unlike in object recognition, where behavioral alignment is often correlated with ImageNet top-1 accuracy, here no such relationship was observed (noise corrected Pearson ρ = 0.09, *p* = 0.87, Fig. S1C). Notably, we observed that models trained specifically on facial identity (VGG-face) or action units (AU: a strategy originally used to generate the emotional intensities in the faces in our imageset^22^) aligned significantly weaker to the macaque behavioral patterns than more broadly trained networks (VGG-Face: 0.43 ± 0.05, AU: 0.44 ± 0.04, Avg. ImageNet Trained ANNs: 0.67 ± 0.04).

Given that static models, despite accuracy matching, failed to fully account for the image-by-image behavioral patterns observed in monkeys, we next examined whether incorporating temporal dynamics into ANN processing could help close this explanatory gap. We evaluated two recurrent and time-extended architectures—CORNet-RT^28^ and ConvRNN^29^—that differ in how they implement temporal processing. CORNet-RT generates internal recurrent dynamics from a static input, whereas ConvRNN processes a temporally structured input sequence matching our stimulus duration. Notably, overall accuracy at each time step was matched to monkey accuracy via unit subsampling, ensuring that any differences reflect representational changes over time rather than differences in ANN performance levels. Interestingly, the two models produced opposite temporal profiles (Fig. 4D). CORNet-RT showed a marked increase in behavioral alignment over time, but doing so in later time windows, suggesting that the additional transformations due to recurrent computations might bring it closer to macaque behavior. In contrast, ConvRNN reached similar alignment earlier but exhibited a significant reduction (Mann-Whitney U test: *U* = 20.0, *n_1_* = 5*, n_2_* = 5, *p* < 0.001) in behavioral consistency with the macaque in later time steps, suggesting that its recurrent processing moves it away from the behavioral signatures observed in macaques. These divergent predictions provide distinct hypotheses for biological data. If IT cortex behaves more like CORNet-RT, alignment with behavior should improve at later stages of processing. If IT behaves more like ConvRNN, alignment should reduce over time. To address this, we performed large-scale neural recordings across the macaque inferior temporal cortex, applying the same behavioral alignment framework to the neural data to test whether IT population activity could predict macaque performance and to evaluate how this relationship evolved over time.

### IT Population Activity Predicts Macaque Behavioral Patterns Better Than Any ANN

We recorded multi-unit spiking activity from 308 neural sites in the inferior temporal (IT) cortex across two macaque monkeys (192 sites in monkey 3, and 116 sites in monkey 4) while they passively viewed the same facial expression images used in the behavioral task (Fig. 5A). We first asked whether distributed patterns of IT activity could discriminate between facial expressions in the human faces shown. We trained a linear classifier on the population response (70-170ms) and estimated decoding performance as a function of expression intensity and category (Fig. 5B). Across expressions, decoding accuracy increased monotonically with intensity—a neural “psychometric” effect mirroring behavior (all slopes > 0; all *p*s < 0.001; Fig. 5B, left). Mean category-level decoding was robust (range: *M* = 0.55–0.70; omnibus ANOVA: *F*(5, 354) = 10.13, *p* < 0.001), with some between-emotion differences (post-hoc, all adjusted *p*s < .05; Fig. 5B, top-right). Critically, these category patterns were reliable across resamples/trials (internal consistency, IC: ≥. 35; Fig. 5B, bottom-right), confirming that the observed neural discriminability reflects stable structure rather than noise. Having established that IT population codes carry sufficient information to discriminate facial expressions, we next asked whether those neural predictions align with the animals’ image-level behavior. Using a time-resolved decoding approach, we trained separate linear classifiers on the population responses within sliding temporal windows and computed performance as decoding accuracy and alignment with macaque behavior. This temporal analysis revealed a distinct temporal dissociation (Fig. 5C): behavioral alignment peaked early (70–100 ms; ρ = .79, *p* < .001) and then declined/plateaued, whereas raw decoding accuracy continued to rise, reaching a later maximum (150–220 ms; *M* = 0.63, Wilcoxon signed-rank test *W* = 51175.0, *n* = 360, *p* < 0.001). Thus, later neural responses contain more categorical information but are less behavior-predictive, indicating that the computations most relevant for decisions are concentrated in an early IT window. To assess sampling limits, we quantified how performance scales with population size (Fig. 5D). Extrapolations indicated that 1297 neurons (early window) or 520 neurons (late window) would be required to reach behavioral-level accuracy (fit early: MSE: 3.6e-07, fit late: MSE: 7.7e-07). Finally, relating neural–ANN representational similarity to behavioral alignment (Fig. 5E) showed a strong association in the late, accuracy-optimal window (noise corrected Pearson ρ = 0.88, *p* < 0.001) but a weaker, nonsignificant trend early (noise corrected Pearson ρ = 0.54, *p* = 0.06), suggesting that ANN–brain similarity does not automatically capture the computations that drive behavior during the behaviorally optimal early phase.

**Fig. 5:**
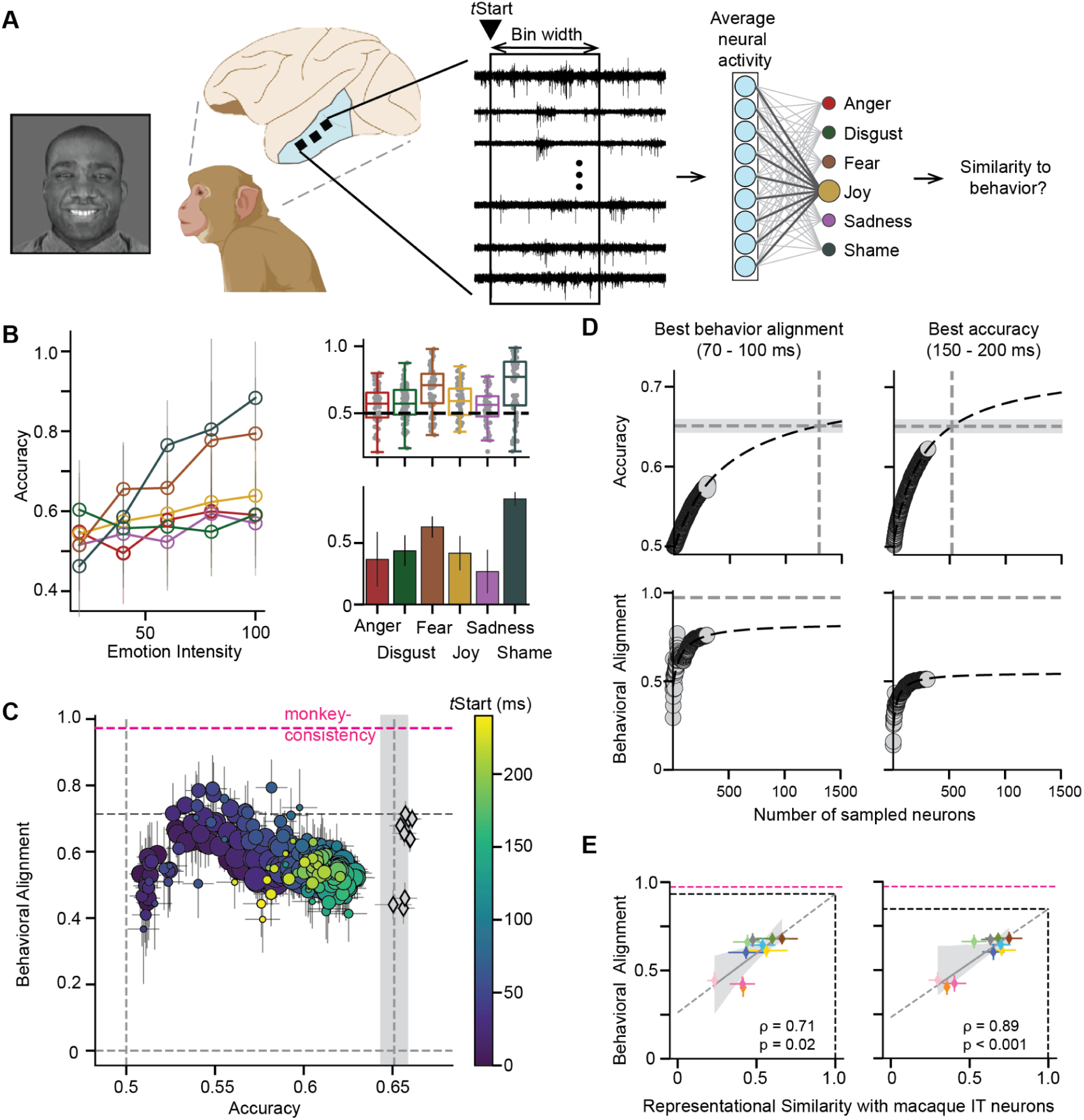
IT population activity predicts macaque facial expression behavior. A) Neural decoding framework. Electrophysiological recordings from inferior temporal (IT) cortex were obtained while macaques passively viewed the same facial expression images used in behavior. Neural spike count was binned in sliding windows, and the time-averaged population activity was used to train six, cross-validated one-vs-rest logistic regression classifiers to predict the displayed emotional category in the image. Classifier predictions were compared to behavioral performance to assess neural–behavioral similarity. **B) Neural decoding performance.** Left: color-coded psychometric function per emotion. Population decoders reliably discriminated facial expressions, with accuracy increasing monotonically with intensity and differing across categories. Right top: Image-level accuracy per expression, with boxes indicating medians and interquartile ranges; each point corresponds to an individual image. Right bottom: Internal consistency (Spearman–Brown corrected split-half reliability) per expression category (mean ± 95% CI). **C) Temporal dissociation between accuracy and behavioral alignment.** Each point corresponds to the decoding accuracy vs. the noise corrected behavioral alignment of the prediction with one time bin (color indicates time from stimulus onset, and size bin width). The black diamonds indicate the model results. The pink horizontal dashed line denotes the monkey-to-monkey consistency and the vertical shaded band highlights the monkey overall accuracy and standard error. While raw decoding accuracy continued to rise late in the response (150–200 ms), behavioral alignment peaked early (70–100 ms), indicating that the computations most relevant for guiding behavior occur in the initial feedforward response. **D) Scaling with population size.**. For the time bin (70-100 ms) with maximal behavioral alignment (left) and maximal accuracy (right) accuracy (top) and behavioral alignment (bottom) are plotted as a function of the number of neurons included. Black dashed lines show extrapolation fits using an arctan. The vertical dashed line indicates where the extrapolation meets the average monkey accuracy (gray horizontal band). Both accuracy and alignment increased with the number of neurons included. Extrapolation revealed that in the early 70–100 ms window, additional neurons would allow performance to reach macaque behavioral accuracy while maintaining the highest behavioral alignment scores, underscoring the behavioral relevance of early IT responses. **E) ANN–neural representational similarity predicts behavioral alignment.** Each dot represents the mean and standard error of the noise corrected representational model-neuron similarity vs. the noise corrected behavioral alignment for each model using the best behavioral alignment bin (left) and best accuracy bin (right). Model colors correspond to Fig. 4. The gray shading denotes 95% CI for the regression line and the black dashed lines indicate the axis interception with 100% similarity. Models whose representational geometry was more similar to IT showed higher behavioral alignment. Both early (behaviorally optimal) and late (accuracy-optimal) windows showed positive relationships, suggesting that model–brain similarity partially explains variance in ANN–behavioral alignment observed in Fig. 4C.

Finally, we asked whether differences in ANN behavioral alignment (as shown in Fig. 4C) could be explained by how well each model’s representational geometry matched that of IT cortex, assessed using representational similarity analysis (RSA). In the early, behaviorally optimal neural window (70–100 ms), when neural activity is most predictive of macaque performance, we found a strong positive relationship between ANN–IT representational similarity and ANN–behavioral alignment (corrected correlation = 0.71; Fig. 5E, left). This suggests that the models whose internal geometry most closely matches IT in this critical early phase are also the ones that better capture the fine-grained behavioral structure of macaques. A similar positive relationship was also observed for the late, accuracy-optimal window (150–200 ms; corrected correlation = 0.89; Fig. 5E, right), indicating that alignment between model and brain geometry is generally beneficial for reproducing behavioral patterns, whether using early decision-related signals or later, more stable categorical representations. Importantly, extrapolating these relationships suggests that if ANNs were to reach the same representational similarity to IT as the monkeys themselves, their behavioral alignment would match the level currently achieved by the neural data. This provides a concrete, quantitative target for ANN development: improving model–brain similarity, particularly in the early decision-relevant phase, could close the current explanatory gap between artificial and biological systems.

### Neural properties and coding strategies underlying facial expression discrimination in IT

The population-level analyses above show that the IT cortex contains rich, temporally dynamic information about facial expression that can predict macaque behavioral patterns. We next asked what neuronal properties best explain this predictive ability, focusing on two candidate factors: (1) a neuron’s selectivity for faces relative to other objects, and (2) its capacity to encode facial identity, and support facial identity discrimination behavior.

We first examined whether a neuron’s selectivity for faces predicts its contribution to facial expression discrimination. Face selectivity was quantified as a Face Selectivity Index (FSI) computed from responses to an independent set of face and non-face object images (Fig. 6A, center). The recorded population spanned a wide range of FSIs, from neurons with no measurable preference for faces (FSI ≈ 0) to neurons responding almost exclusively to faces (FSI > 0.4). Example tuning profiles illustrate that intermediate-FSI neurons responded robustly to both faces and non-faces, whereas highly face-selective neurons responded almost exclusively to faces (Fig. 6A, left and right). To relate face selectivity to expression discrimination, we examined how each neuron’s FSI value correlated with its absolute regression coefficient from the most behaviorally aligned facial expression decoder, computed separately for each expression category (Fig. 6B). Across all six expressions, we observed a consistent positive relationship between face selectivity and decoder weight (*r* = 0.17–0.37, all *p* < 0.05), indicating that more face-selective neurons generally contributed more strongly to expression decoding. This relationship was stable across emotions, suggesting that face selectivity is a general predictor of a neuron’s importance for expression discrimination.

**Fig. 6:**
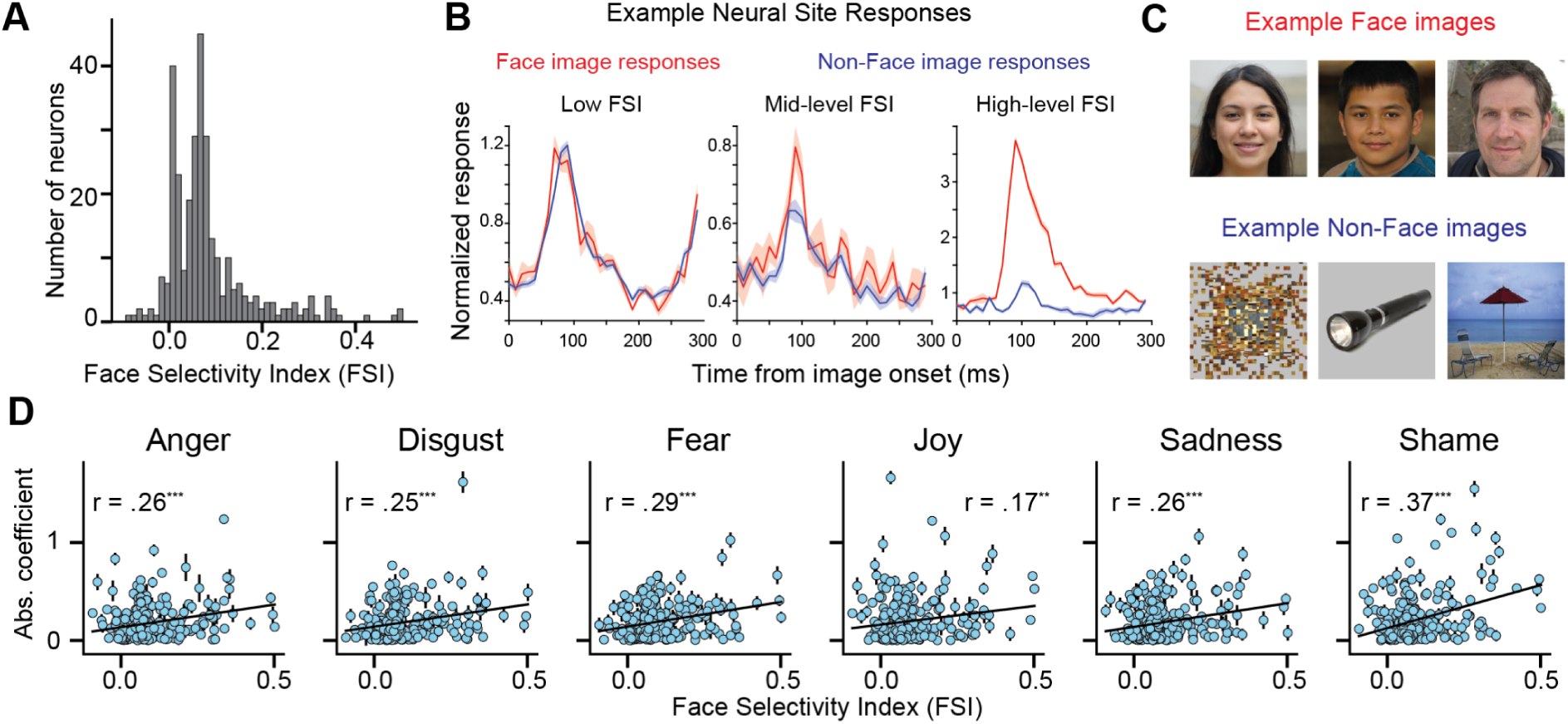
Face selectivity partially predicts neuronal contributions to facial expression discrimination. A) Distribution of face selectivity indices in our recorded neural population. The histogram shows the distribution of Face Selectivity Index (FSI) values across all recorded sites. **B) Example responses of neurons with different degrees of face-selectivity.** Example neural responses to face images (red) and non-face (blue) images for sites with low FSI (left), with intermediate FSI (middle), and with strong FSI (right). **C)** Example face and non-face images used in the study to estimate the face selectivity indices are shown on the far right. To ensure anonymity, we replaced all face images with generated images.^30^ **D) Expression decoding strength increases with face selectivity.** Across all six expressions, neurons with higher FSI values tended to exhibit larger regression coefficients in the expression decoder, indicating stronger—but not exclusive—contributions to discrimination performance. For each expression category, scatter plots show the relationship between FSI and the absolute value of the corresponding logistic regression coefficient (averaged across cross-validation folds), reflecting the contribution of each neuron to predicting that expression. The error bars indicate the standard error across cross-validation folds. The black lines show least-squares fits. Pearson’s *r* values (0.17–0.37) and significance levels are shown for each category (p < 0.01 = **; p < 0.001 = ***).

We next asked whether the same neurons that carry strong expression information also support facial identity discrimination. Using a separate identity discrimination task with the same face stimuli (Fig. 7A), we found that many neurons achieved high identity decoding accuracy (Fig. 7B, *M*=0.90, Wilcoxon Signed-Rank test: *W* = 64620, *n* = 360, *p* < 0.001). Comparing behavioral alignment across the two tasks revealed a strong positive correlation (noise corrected Pearson ρ = 0.85, *p* < 0.001; Fig. 7C): neurons whose responses were better aligned with behavior in the identity task were also better aligned in the expression task. This relationship persisted when comparing average decoder coefficients (noise corrected Pearson ρ = 0.76, *p* < 0.001; Fig. 7D) and when comparing raw decoding accuracies (noise corrected Pearson ρ = 0.50, *p* < 0.001; Fig. 7E). Color-coding neurons by FSI in these analyses revealed that many of the strongest dual-task performers fell within the mid-to-high FSI range identified in Fig. 6. Importantly, we did not observe a significant correlation between the facial identity and expression discrimination behavioral patterns for all emotion categories except shame (anger: ρ = 0.01, *p* = 0.92; disgust: ρ = 0.21, *p* = 0.12; fear: ρ = -0.06, *p* = 0.66; joy: ρ = -0.02, *p* = 0.87; sadness: ρ = 0.11, *p* = 0.39; shame: ρ = -0.62, *p* < 0.001; all ρ are noise corrected). To test whether the strong negative correlation observed for shame biased the relationship between neural decoder alignment for identity and expression, we recomputed the analysis after excluding shame images. The correlation remained significant (Pearson *r* = 0.54, *p* < 0.001), confirming that behavioral similarity across tasks did not drive neural decoder alignment. This reinforces the conclusion that expression discrimination emerges from a partially overlapping but functionally distinct subspace, rather than from identical identity-related codes. Together, these results indicate that the neurons most predictive of behavioral performance in facial expression discrimination are often both highly face-selective and identity-informative. This convergence points to a shared, partially overlapping representational subspace (and likely readout differently in the downstream areas) in IT cortex that supports both identity and expression recognition—a subpopulation that may form the core neural machinery for decoding socially relevant facial information.

**Fig. 7:**
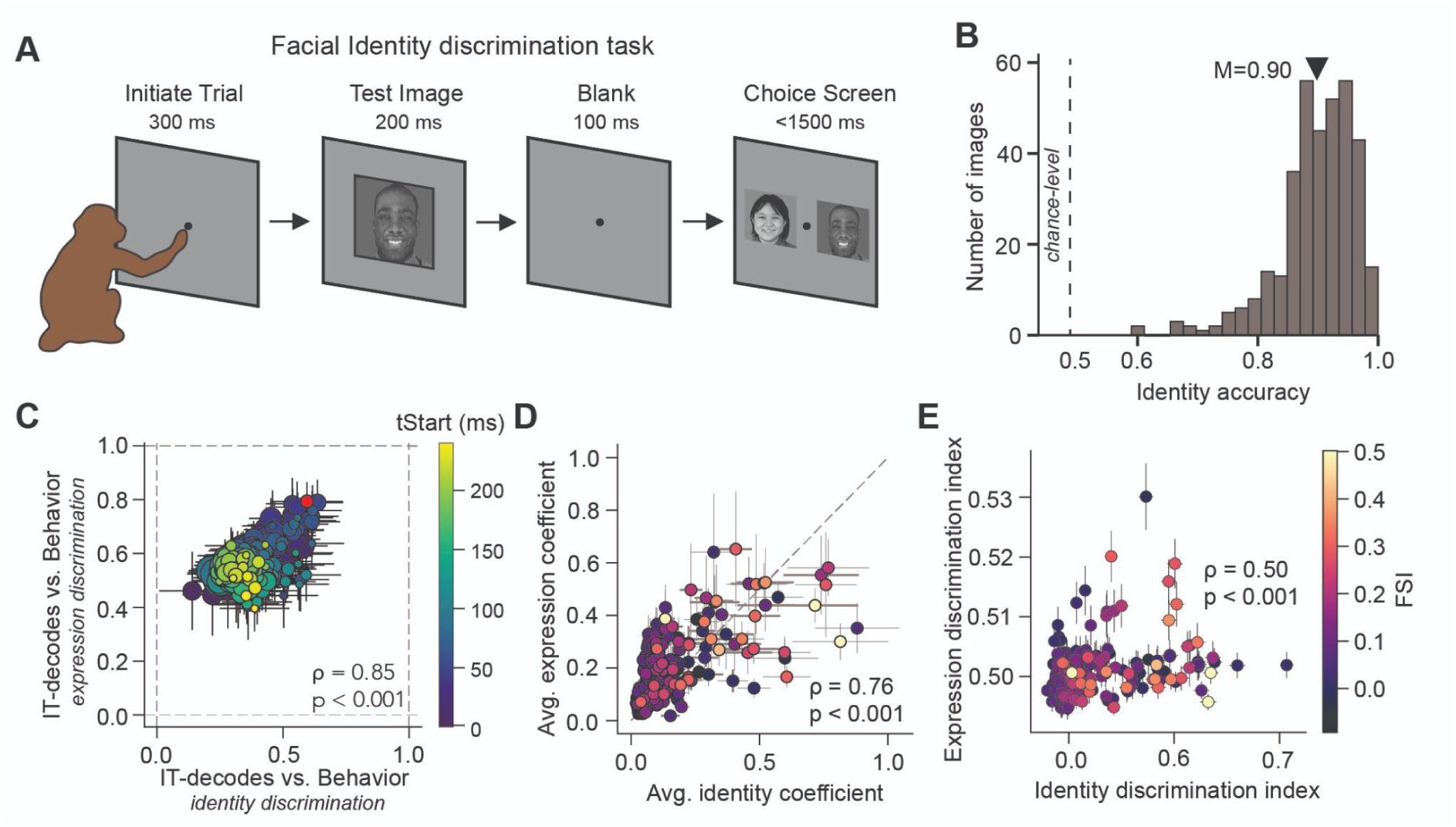
Overlap between identity and expression coding in macaque IT neurons. A) Schematic of the identity discrimination task. After monkeys initialize a trial by tapping the screen, a Test image is presented for 200 ms, followed by a 100 ms delay. Monkeys then choose the identity of the Test image from a forced choice between the correct identity (target) and a distractor (one of the other eleven identities), both shown with the same expression at 100% intensity. **B) Distribution of macaque behavioral accuracies for identity discrimination.** The population mean is indicated by the black triangle (*M*=0.90). **C) Comparison of IT population decode vs. behavior alignment for identity discrimination and expression discrimination.** Each dot represents one time window and the color and size indicate the start time (tStart) and bin width used to develop the IT decoder, respectively and the error bars show the standard deviation across 100 bootstrapped samples across images. The red dot refers to the highest expression behavior alignment decoder choice (built from activity binned between 70 to 100 ms post image onset). This was used for subsequent analyses in panels D and E. **D) Strong overlap between individual neuron’s contribution to facial identity and expression discrimination.** Single neural sites that contributed strongly to identity discrimination also tended to contribute strongly to expression discrimination (corrected Pearson correlation = 0.76, p<0.001). Each dot and error bar represent the mean and standard error across the absolute coefficient for all identities or expressions. The dashed line shows the bisector. **E) High correlation between single neuron’s identity and expression discrimination indices.** Neurons that achieved high identity decoding accuracy also tended to achieve high expression decoding accuracy (corrected Pearson correlation = 0.5, p<0.001). For each neuron we trained a one-vs-rest logistic regression classifier to predict identity and expression. Each dot and error bar shows the average and standard error of the image-level prediction performance per neuron. In (D) and (E) all dots are color-coded by FSI.

## Discussions

Our study provides a unified, multi-level analysis of facial expression discrimination, connecting human and macaque behavior, macaque IT neural activity, and ANN models within the same image-level framework. By aligning these domains at high resolution, we identify candidate mechanistic features of how identity and expression are represented in the primate visual system and set quantitative targets for building more brain-like computational models.

### Macaques as a high-fidelity model for human facial expression discrimination

We demonstrate that macaques not only learn a human facial expression discrimination task to high accuracy (Fig. 3A-B) but also exhibit image-by-image performance patterns that closely parallel human observers across all six expressions (Fig. E-F). This strong cross-species alignment—comparable to that reported in object recognition^26^—validates macaques as a model for probing the sensory aspects of the neural basis of human facial expression processing. It also justifies the use of human face stimuli in macaque neurophysiology, enabling direct tests of human-relevant hypotheses about the neural computations underlying social perception. From a broader perspective, this result addresses a long-standing gap in social vision research: while macaques are a well-established model for face processing, rigorous image-level behavioral validation of their suitability for human facial expression studies has been lacking. By providing this benchmark, we establish a foundation for mechanistic experiments and causal interventions whose inferences can be potentially translated to human social cognition.

### ANNs reveal representational gaps and divergent temporal predictions

Evaluating a diverse set of ANN models, we find that decoders built from activations in the later, IT-like layers better align with macaque behavioral patterns than earlier layers, consistent with the hierarchical nature of ventral stream processing (Fig. 4B), and with the idea that higher stages carry behaviorally relevant invariances^31^. Critically, even after *accuracy matching* via unit subsampling (controlling for performance confounds), no static model—including those explicitly trained on facial identity or action units (AU)^19^—fully captures macaque behavior (Fig. 4C), underscoring a representational gap between current ANNs and primate expression discrimination. This result is in line with recent AU-model evaluations showing that AU-based hypotheses explain *some* variance in human expression judgments but remain well below noise ceilings, indicating missing features and constraints in the current modeling space^21^. Likewise, work on identity-trained face ANNs demonstrates that expression-selective units *can* emerge and capture aspects of human-like expression perception, but only under domain-specific face experience—and still with accuracy shortfalls relative to humans—again pointing to incomplete mechanisms in off-the-shelf training regimes^32^. Consistent with these observations, we observed a lack of correlation between ImageNet accuracy and behavioral alignment reinforcing that expression discrimination might depend on representational axes not emphasized or evolved by the objectives of object categorization in ANNs. Beyond static models, our time-resolved tests reveal that recurrent architectures make *opposing, falsifiable* predictions about neural dynamics: CORNet-RT’s^28^ behavioral alignment increases over time, whereas ConvRNN’s^29^ alignment declines at later time bins—even when accuracy is matched at each step—implying divergent recurrent transformations over the same inputs (Fig. 4D). This divergence positions ANN modeling not merely as a descriptive tool but as a generator of specific, falsifiable predictions about neural computations across time.

### Early IT responses are most behaviorally aligned

Direct IT recordings revealed a temporal dissociation between *encoding* and *behavioral relevance*. Behavioral alignment peaked early, within 70–100 ms of stimulus onset, whereas raw neural decoding accuracy continued to rise, reaching its maximum later (150–200 ms). This indicates that the computations most relevant for guiding expression judgments occur during the initial feedforward sweep of IT activity, consistent with prior object recognition studies showing that early IT signals are sufficient to drive core recognition behavior^33^. Later IT responses, although more categorical, may reflect recurrent refinement, feedback from higher-order regions, or post-decisional processes not directly engaged by the task. This dissociation underscores a broader principle: encoding capacity and behaviorally deployed information are not synonymous, and early visual signals often play the decisive role in perceptual judgments. Our findings also converge with recent computational work demonstrating that activations of feedforward models can explain the behavioral variance predicted by human amygdala responses^34^. Taken together, these studies suggest that early ventral-stream representations are sufficient to support behaviorally relevant judgments of facial expressions. Our neural dynamics further adjudicate the opposing predictions made by recurrent ANNs. CORNet-RT^28^ exhibited increasing behavioral alignment over time, while ConvRNN^29^ alignment declined in later time bins, despite both models being accuracy-matched at each time step. The IT data align more closely with the ConvRNN profile, where later recurrent activity does not add, but rather deteriorates, behaviorally relevant structure, providing a rare instance where ANN dynamics generate falsifiable hypotheses that are directly tested in the brain. At the same time, previous work has established the importance of late-phase IT activity for object recognition under challenging conditions, where recurrent processing helps to build more accurate codes^17,35^. A similar principle may hold for faces: while early IT signals dominate in standard discrimination tasks, late activity could become critical for coding expressions in more ambiguous or noisy contexts. Thus, we speculate that the current gap between neural predictivity and the behavioral ceiling may be closed by developing more accurate dynamic decoding models that optimally integrate early feedforward signals with recurrently refined late responses.

Finally, relating ANN–brain representational similarity (RSA) to ANN–behavior alignment revealed that models with greater IT-like representational geometry also achieved stronger behavioral alignment, particularly in the late window. Importantly, extrapolation suggests that if ANNs were trained to achieve monkey-level RSA^36^, they would also reach neural-to-behavior alignment. This defines a quantitative and biologically grounded target for future ANN development, moving beyond generic “brain-likeness” toward measurable alignment with both neural activity and behavior.

### Face selectivity supports expression discrimination

At the single-neuron level, we find a robust positive relationship between face selectivity (FSI) and contribution to expression decoding across all expressions. This indicates that, on average, more face-selective neurons tend to contribute more strongly to expression discrimination, although some less-selective neurons also carried substantial weights. Thus, while face selectivity is not an exclusive requirement for expression coding, but it biases neurons toward stronger contributions. This dependency on face selectivity also indicates a hierarchical relationship between structural and dynamic aspects of face processing—identity-stable structural encoding appears to form the scaffold upon which expression discrimination is built. In computational terms, neurons tuned to the high-dimensional subspace of face identity may be best positioned to capture within-identity variations corresponding to expressions, but our data also highlight heterogeneity in how neurons across the FSI spectrum contribute.

### Identity–expression overlap reveals a shared representational subspace

Our analyses reveal that facial expression discrimination in IT cortex is not the sole function of a specialized, expression-only circuit, but instead emerges from a subpopulation of neurons that also carries substantial facial identity information. These neurons, typically in the mid-to-high range of face selectivity, form a shared representational substrate in which identity and expression are encoded along partially overlapping dimensions. This architecture challenges the long-standing view that the most face-selective neurons are the sole repository of behaviorally relevant expression information. Instead, our data show that identity- and expression-relevant codes coexist within the same subspace, but are weighted differently depending on task demands. This overlap may provide a computational advantage: by maintaining stable structural features for identity while preserving sensitivity to changeable cues, the system can flexibly support both social recognition and social communication. Importantly, this framework generates a testable prediction—perturbing these identity-informative, face-selective neurons should produce coupled deficits in both identity and expression discrimination. More broadly, our findings suggest that the principle of shared but differentially weighted subspaces may generalize to other domains in high-level vision, offering a unified model for how IT supports multiple behaviorally critical functions within a common representational geometry.

### Limitations

Despite the strengths of our integrative approach, several limitations remain. While macaques and humans perform the task similarly, behavioral discrepancies between the species remain, particularly for *Disgust* and *Anger*, highlighting species-specific strategies that may constrain generalization (Fig. 3E). Moreover, a gap remains between prediction and behavior, not only for artificial models, but also for the best-predicting neurons in IT cortex, suggesting that additional factors beyond local visual representations likely contribute to behavioral performance. Our analyses address only the visual encoding of facial expressions, however, we cannot capture how feedback from higher-order or affective brain regions may modulate these representations to infer the emotional meaning in humans. While we observed that information in IT unfolds dynamically, our current analyses did not explicitly exploit these temporal patterns, leaving open the possibility that richer dynamic signals may further improve predictions. Finally, although facial expressions are dynamic signals, we focused on static images to provide precise experimental control. Future studies incorporating videos or morph sequences could better capture the temporal structure of expression processing.

### Broader implications

Taken together, our results support a model in which early IT responses from a mid-to-high FSI, identity-informative subpopulation form the core substrate for both identity and expression recognition. This shared representational subspace integrates stable and dynamic facial features, providing a flexible platform for multiple social vision tasks.

From a translational perspective, this framework also offers a mechanistic entry point into disorders of social perception. Differences in facial expression discrimination are a hallmark of autism, with autistic adults often showing atypical image-level patterns compared to neurotypical individuals^37,34^. By establishing macaques as reliable models of human facial expression recognition and aligning behavior, IT neural activity, and ANN predictions at the image level, our study provides a circuit-level platform for modeling social perception differences in autism. This framework enables ANN-based predictions about how neural coding strategies might differ in autism — for example, altered weighting of identity vs. expression dimensions — and provides a primate system in which such hypotheses can be causally tested.

From a modeling standpoint, our ANN analyses reveal both representational gaps and explicit engineering targets: models that better match IT representational geometry, especially in the early behaviorally optimal phase, should yield improved alignment with behavioral structure. From a neuroscience perspective, this framework unifies identity and expression coding, reframing them not as segregated streams but as intertwined dimensions of a common code.

Finally, the cross-species image-level alignment we establish enables causal testing of these principles: perturbations targeting mid-to-high FSI, identity-informative neurons should jointly impair identity and expression discrimination in predictable ways. More broadly, the principle of shared but differentially weighted subspaces may generalize to other high-level visual domains, offering a unified account of how IT supports multiple behaviorally critical functions within a common representational geometry.

In sum, by integrating cross-species behavior, direct neural recordings, and model-based predictions, we move beyond descriptive accounts to a mechanistic framework that unites identity and expression coding in IT, identifies the subpopulation most critical for social vision, and defines quantitative targets for building models that truly match brain and behavior.

## Methods

### Visual Stimuli

We used a combination of grayscale photographs from the Montreal Set of Facial Displays of Emotion (MSFDE) dataset.^22^ The dataset includes six expression categories: angry, disgust, joy, fear, sadness, and shame. We selected a subset of twelve identities (6 female), four individuals of each ethnicity (European, Asian, and African). Each individual was photographed while expressing an emotion, resulting in photographs with 20-100% of the specific emotion displayed. In each image, we equalized low-level properties using the SHINE (spectrum, histogram, and intensity normalization and equalization) toolbox.^3^

We used a separate set of 60 images to determine the face selectivity index of the neural sites. This imageset contained 15 static faces, 15 objects, 15 scenes, and 15 scrambled objects.

### Behavioral training and testing

#### Human participants

We collected large-scale psychophysical data from 290 subjects using Amazon Mechanical Turk (MTurk). Participants completed the tasks on MTurk, an online crowdsourcing platform, for a payment of $15 CAD/hour. This experimental protocol involving human participants was approved by and in concordance with the guidelines of the York University Human Participants Review Subcommittee. Humans did not receive any additional training on the task. Each trial began with a brief presentation (200 ms) of a sample image with a given emotion, selected from the set of 360 images. After a 100 ms blank gray screen, subjects were shown a choice screen displaying an image of a target and a distractor emotion. Subjects indicated their choice by clicking on the image they believed matched the emotion of the sample image. To ensure high-quality behavioral data, we rejected all participants who performed below an accuracy of 0.6 (chance-level = 0.5) on the first 30 trials during each session. Furthermore, we evaluated the reliability of the image-level behavioral metrics as a function of the number of trials (Supplementary Fig. S2A) and found that the reliability of the mean image-level accuracy pattern increased with more repetitions per image and reached high values (∼0.8), nearly asymptoting with around 60 trials.

#### Non-human primates

We trained two adult male rhesus macaque monkeys (Macaca mulatta; monkey 1 and 2) to perform the emotional expression and identity discrimination task. All data were collected, and animal procedures were performed, in accordance with the NIH guidelines, the Massachusetts Institute of Technology Committee on Animal Care, and the guidelines of the Canadian Council on Animal Care on the use of laboratory animals and were also approved by the York University Animal Care Committee.

The two rhesus macaques were trained to perform the two-alternative forced-choice (2AFC) facial expression discrimination task using a match-to-sample paradigm.^39^ Training was organized into sequential phases of increasing difficulty, gradually introducing additional emotion categories, identities, and subtler intensity levels. Monkeys trained 5 days per week in sessions of ∼1000–2000 trials, progressing to the next phase upon reaching ≥80% correct performance over two consecutive sessions. Each trial began with fixation on a central dot, followed by a 500ms presentation of a sample face (neutral or emotional; ∼8° visual angle), a brief 100ms blank delay, and a choice screen displaying two faces of the same identity: one matching the sample’s emotion category at full intensity and one with a distractor expression. In early training stages, the correct choice was identical to the sample image to facilitate learning. Monkeys indicated their choice by touching one of the two images, with correct responses rewarded by water and incorrect responses followed by a short timeout (5s) before the next trial. This positive reinforcement protocol was maintained throughout training to promote stable and high-accuracy performance.

During behavioral testing images were presented on an iPad^25^. All images were shown at 8 deg of visual angle. Monkeys touched the screen to initiate a trial. Similar to the human participants, the trial started with the presentation of a sample image from the set of 360 images for 200 ms. This was followed by a blank gray screen for 100 ms, after which the choice screen was shown displaying a target and a distractor image. For the expression discrimination task, the identity was fixed, while for the identity discrimination task, the shown emotion was fixed and displayed at 100%. The monkey was allowed to view the choice screen freely for up to 1500 ms and indicated its final choice by touching the selected image.

Using this set up, monkeys performed both the expression and the identity discrimination task. Prior to testing in the laboratory, monkeys were trained to perform both discrimination tasks.

#### Behavioral metrics and behavioral alignment

We developed a behavioral metric to quantify the performance in the discrimination task on the level of individual images. This image-level accuracy metric (B.I1, see Fig. 1) estimates the discriminability of each image, pooling across all distractors. Given our image set of 360 images, B.I1 comprises 360 values. Specifically, given an image *i* of a given category *c* ∈ *C* and all distractor categories (*d* ∈ {1, ···, *D*} and *c* ∉ *D*), where *C* and *D* are both either expression or identity specific, we computed the average performance per image as:

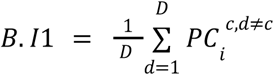

where 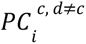 indicates the percent correct for a given image *i* as the fraction of correct responses in the binary task, between a category *c* and a distractor *d* ∈ *D*.

We quantified the internal consistency of B.I1 using Spearman–Brown–corrected split-half reliability. For a given monkey, all trials for each image were randomly divided into two equal halves across *N* iterations (here, *N*=100). The behavioral metric was computed separately for each half, and the Pearson correlation (*r_half_*) between the two halves was calculated across images. The mean correlation across all iterations was then adjusted using the Spearman–Brown correction to account for the reduced number of trials in each half, providing an estimate of the reliability for the full dataset:

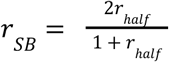

All reported “noise-corrected” correlations involving behavioral data were normalized by the square root of the product of the relevant internal consistencies.

To assess behavioral alignment, we computed the correlation between the model- or neuron-derived image-level predictions (B.I1) and the corresponding macaque B.I1. To account for measurement variability, we corrected the raw correlations using the geometric mean of the internal consistencies of the correlated values:

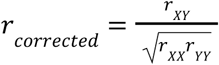

where *r_XY_* is the observed correlation, and *r_XX_*, *r_YY_* are the internal consistencies of the neural predictions and behavioral metrics, respectively. As the tested models are not stochastic by nature, we assumed *r_XX_*= 1 for all model predictions

### Artificial neural networks (ANNs)

#### Generating behavioral predictions from ANN activations

We first evaluated ANNs to build hypotheses about the underlying mechanisms in macaque facial expression discrimination. We therefore evaluated nine static (time-agnostic) and two dynamic (time-recursive) convolutional neural networks (CNNs, see Table 1 for model characteristics and training objectives). These models included a diverse set of static models such as standard architectures (AlexNet, ResNet18, and ResNet50), contrastive learning models with and without language input (CLIP^40^ and SimCLR,^41^ respectively), a model trained to predict facial Action Units (AU), and CORNet-Z^28^ as representative of biologically inspired models. We also included domain-specific models trained exclusively on human faces to perform an identity discrimination task or exclusively on objects to perform object categorisation.^42^ In addition, we analyzed dynamic models, including CORNet-RT^28^, and ConvRNNs,^29^ that incorporate recurrent processing to approximate temporal dynamics more closely aligned with biological systems. For each model, we extracted the unit activations from each layer in the models for every stimulus. We mimicked the macaque expression discrimination task by training six one-vs-rest logistic regression classifiers on the activations from each model layer (Fig. 4A) using 5-fold stratified cross-validation At the conclusion of ANN fine tuning for expression prediction, we would generate 6 probabilities (one for each emotion) when a Test image was presented to the ANN.

**Table 1:** Details of the ANN models used in this study.

| Model name | Model category | Training data | Training objective | IT layer |
| --- | --- | --- | --- | --- |
| AlexNet<sup>43</sup> | Generic CNN | ImageNet | Object classification | features.12 |
| ResNet18<sup>44</sup> | Residual CNN | ImageNet | Object classification | layer4.0.relu |
| ResNet50 | Residual CNN | ImageNet | Object classification | layer4.1.relu |
| CLIP<sup>40</sup> | Contrastive | Images & Text | Self-supervised | visual.layer4.0.relu2 |
| SimCLR<sup>41</sup> | Contrastive | ImageNet | Self-supervised | net.4.blocks.1.net.1.1 |
| CORNet-Z<sup>28</sup> | Biologically inspired | ImageNet | Object classification | IT.output |
| ME-GraphAU<sup>45</sup> | Facial action units | BP4D<sup>46</sup> | Facial action units | output |
| VGG-Face<sup>18</sup> | Face-trained | Human faces | Identity classification | fc7 |
| VGG-Object<sup>18</sup> | Object-trained | Objects | Object classification | fc7 |
| CORNet-RT<sup>28</sup> | Recurrent | ImageNet | Object classification | IT.output |
| ConvRNN<sup>29</sup> | Recurrent | ImageNet | Object classification | conv_7 |

#### Generating task accuracies from output probabilities

To generate a comparable trial-averaged performance per image similar to the monkeys and

humans, a probability for each classifier output given any image *i* of category *c* (*P_i_*^*c*^) was generated.

We then computed the binary task performances by calculating the percentage correct score for each pair of possible binary tasks given an image. For instance, if an image was from category *c*, then the percentage correct score (PC) for the binary task between the target category *c* and the distractor category d, *PC_i_*^*c*,*d*^ was computed as follows:

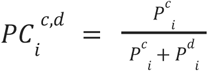

From each percentage correct score, we then estimated an ANN *I*1 score (per image), following the same procedures as the behavioral metric. To enable a fair comparison with macaque behavior, we iteratively subsampled units from each layer until the classifier’s accuracy matched that of the monkeys. This subsampling was implemented via a binary search procedure, in which a random subset of units was drawn, its performance computed, and the process repeated until the target accuracy was reached. The reported results reflect the average prediction accuracy and behavioral alignment across 50 iterations of the matched subsets.

#### Selection of the ANN-IT layers

To ensure that our ANN–neural comparisons are anchored at the most biologically relevant processing stage, we selected for each model the layer with the highest alignment to macaque IT cortex as determined by the BrainScore benchmark. This approach maximizes the interpretability of behavioral alignment results, as it isolates the stage in the model most likely to capture IT-like representational structure. For the Action Unit (AU) model, we instead used only the AU-based predictions to directly assess the explanatory power of action units for macaque behavior. For the face- and object-trained models, we used the final classification layer, which should, by design, have the highest discriminatory power for the categories on which the model was trained (identities for face-trained, object classes for object-trained). This choice allows us to test whether a model’s most category-selective stage, where task-relevant features are most strongly amplified, also best aligns with the behavioral patterns observed in macaques. For dynamic models, we treated each time step as an independent representational state, training a classifier at each time step and subsampling units to match macaque accuracy. This procedure allowed us to track the temporal evolution of ANN–behavior alignment and compare it directly to the dynamics observed in neural data.

#### Early vs. late processing stages

To determine whether early or late processing stages better align with macaque behavior, we analyzed the first 30% (early) and last 30% (late) of layers from each static model (Fig. 4B). The action unit model was excluded from this comparison, as we only collected the general AU predictions, as were the face- and object-trained models, where only the last fully connected layers were available. For each selected layer, we computed the behavioral alignment between macaque *I*1 and the accuracy-matched model *I*1 predictions. We then averaged these alignment values within the early and late layer groups, and report the mean and standard error of these averages across models.

##### Electrophysiological recording and data preprocessing

We recorded neural activity to the subset of 360 images of the MSFDE imageset, and the 60 images from the Face Object Scene Scrambled (FOSS) imageset. Each image was resized to 256 x 256 x 3 pixel size and presented within the central 8 deg.

In addition to the two behaviorally trained monkeys, we had two different male rhesus monkeys (Macaca mulatta; monkey 3 and 4) that were surgically implanted with a head post under aseptic conditions. All data were collected, and animal procedures were performed, in accordance with the NIH guidelines, the Massachusetts Institute of Technology Committee on Animal Care, and the guidelines of the Canadian Council on Animal Care on the use of laboratory animals and were also approved by the York University Animal Care Committee. Neural activity was recorded using two or three micro-electrode arrays (Utah arrays; Blackrock Microsystems) implanted in IT cortex (see Fig. S3). A total of 96 electrodes were connected per array (grid arrangement, 400 µm spacing, 4mm x 4mm span of each array). Array placement was guided by the sulcus pattern, which was visible during the surgery. The electrodes were accessed through a percutaneous connector that allowed simultaneous recording from all 96 electrodes from each array.

During each recording session, band-pass filtered (0.1 Hz to 10 kHz) neural activity was recorded continuously at a sampling rate of 20 kHz using Intan Recording Controllers (Intan Technologies, LLC). The majority of the data presented here were based on multiunit activity. We detected the multiunit spikes after the raw voltage data were collected. A multiunit spike event was defined as the threshold crossing when voltage (falling edge) deviated by less than three times the standard deviation of the raw voltage values. Our array placements allowed us to sample neural sites from different parts of IT, along the posterior to anterior axis. However, for all the analyses, we did not consider the specific spatial location of the site, and treated each site as a random sample from a pooled IT population.

All neural response data were obtained via a passive viewing task. In this task, monkeys fixated a white square dot (0.2°) for 300 ms to initiate a trial. We then presented a sequence of 5 to 10 images, each ON for 100 ms followed by a 100 ms gray blank screen. This was followed by a water reward and an inter trial interval of 500 ms, followed by the next sequence. Trials were aborted if gaze was not held within ±2° of the central fixation dot during any point. Firing rates were computed using a non-overlapping sliding window of 10 ms to capture the temporal dynamics of the neural responses.

#### Exclusion criteria

We only included neural sites with high internal consistency in our analyses. Specifically, we computed the Spearman-Brown corrected split-half reliability for each neuron across 100 random half-splits across all trials:

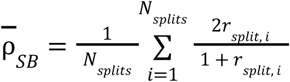

where *N_splits_* is the number of random half-splits, and *r_split_* the Pearson correlation between the responses in two random halves of the trials for that neuron. Neural sites were excluded if their split-half consistency, measured over a time window of 70–170 ms post-stimulus onset, fell below 0.3. This window corresponds to the typical visually evoked response latency in IT cortex and has been shown to correlate with object classification behavior in primates. Following this quality control step, the final dataset included 308 neural sites (192 from monkey 3 and 116 from monkey 4).

#### Generating behavioral predictions from neural data

To ask how a downstream readout region might utilize the temporal dynamics of the IT cortex to support facial expression and identity discrimination,, we modeled the binary discrimination task using one-vs-rest logistic regression classifiers (Fig. 5). Each classifier was trained with 5-fold stratified cross-validation. We generated ∼200 independent decoders based on a combination of decode start time and integration time bin to evaluate how discriminability varied over time. Each neural decoder was trained to generate 6 expression probabilities or 12 identity probabilities when shown an image at the test phase. We then convert these probabilities into neural population *I*1 image-level vectors following the same procedures as reported above for the ANNs

#### Sampling analysis

To assess how the neural prediction’s accuracy and correlation with behavior scales with population size, we repeated the decoding procedure while systematically subsampling the number of neurons included in the decoder (Fig. 5D). Specifically, we randomly sampled an increasing number of neurons without replacement, ranging from a small subset up to the full recorded population. For each sampling size, we trained the classifier and computed the corresponding performance metric, repeating the process over 100 random draws to obtain a stable mean estimate. The resulting mean performance (accuracy or behavioral alignment) as a function of sampled neuron count was then fit with an arctangent:

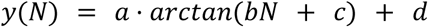

where *N* denotes the number of sampled neurons, *a* controls the amplitude, *b* represents the growth rate, *c* indicates the horizontal shift and *d* is an offset term. This parametric fit allowed us to extrapolate performance beyond the recorded population size and estimate the number of neurons required to reach behavioral performance levels observed in the macaques.

#### Predictions from single neurons

To assess the contribution of individual neurons to expression or identity decoding, we repeated the prediction procedure using the activity of single neurons in isolation (Fig. 7E). We refer to them as the *expression discrimination index* and the *identity discrimination index* for each neuron (Fig 7E). For each neuron, we averaged the response within a specified temporal window, and trained a one-vs-rest logistic regression classifier to predict facial expressions or identities. Therefore for each image, we generate 6 probabilities for expressions and 12 probabilities for the identities. We then convert these probabilities into neural *I*1 image-level vectors following the same procedures as reported above for the ANNs and neural population decoders. Classification accuracy was evaluated using 5-fold stratified cross-validation. We report the resulting average *I*1 accuracy for each neuron as the corresponding expression or identity discrimination index, enabling us to identify units with above-chance predictive power and to examine how single-neuron encoding dynamics contribute to population-level performance.

##### Representational Similarity Analysis

To test whether models that share greater representational similarity with IT neural responses also achieve higher behavioral alignment, we performed a representational similarity analysis (RSA)^47^ between model features and neural population responses (Fig. 5E). For each model, we extracted the same feature set used in the behavioral prediction analysis, i.e., unit activations subsampled to match macaque-level prediction accuracy. For the neural data, we averaged the neural response within a predefined temporal window of interest. We then computed a representational similarity matrix (RSM) by calculating the Pearson correlation of all features or neurons across all 360 images. Model–brain similarity was then quantified by correlating the flattened model and neural RSMs. To account for measurement noise, the similarity values were corrected by dividing them by the square root of the split-half reliabilities of the neural RSMs estimated across trials for each neuron using Spearman–Brown correction.

##### Face selectivity index (face vs. non-face)

We computed the face selectivity of each neuron to test if expression- or identity-predictive neurons are also strongly face-selective. We quantified the face selectivity of each neuron using a specific dataset that contained different images of objects, scenes, scrambled images, as well as faces. Each image was resized to 256 x 256 x 3 pixel size and presented within the central 8 deg to the monkey.

For each recorded neuron, we computed the average response to face images *r_F_* and to non-face images *r_NF_*, and calculated a face selectivity index (FSI) as:

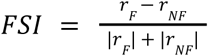

This index ranges from -1 to 1, where positive values indicate a preference for faces, negative values indicate preference for non-face stimuli, and values near zero suggest no specific selectivity.

## Acknowledgments

KK has been supported by funds from the Canada Research Chair Program, the Simons Foundation Autism Research Initiative (SFARI, 967073), Brain-Canada Foundation, the Canada First Research Excellence Funds (VISTA Program), and the National Sciences and Engineering Research Council of Canada (NSERC). MW received funding from the Connected Minds Postdoctoral Fellowship (supported by CFREF), the Alfons and Gertrud Kassel-Stiftung and the Deutsche Forschungsgemeinschaft (DFG, German Research Foundation) – 414985841. HR was supported by a CIHR Postdoctoral Fellowship.

## Supplementary Material

**Fig. S1:**
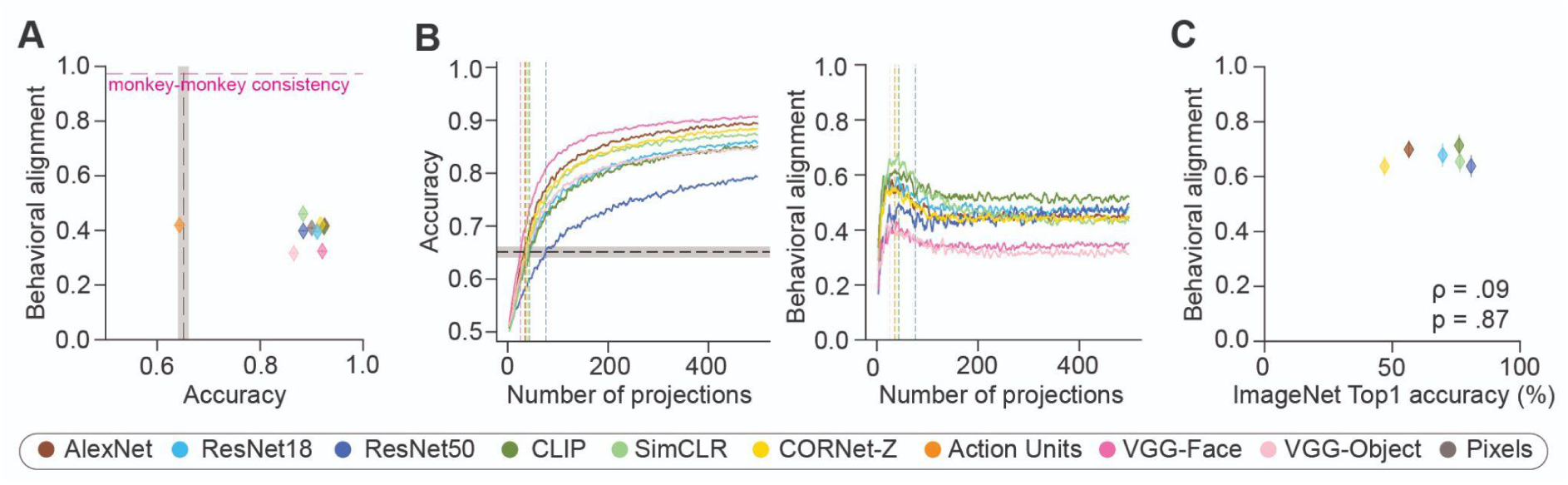
Artificial neural network performance. **A)** Random projections of the full IT layer (500 dimensions) reveal that higher classification accuracy does not necessarily lead to better behavioral alignment with macaques. Each datapoint represents the prediction accuracy vs. the noise corrected behavioral alignment of one ANN model. The gray translucent band shows the average pooled monkey accuracy and the pink horizontal line indicates the consistency of the image-level accuracy patterns between the two monkeys. **B)** Sampling analysis from model IT layers. Left: increasing the number of sampled units improves accuracy. Right: behavioral alignment initially increases but declines at higher accuracies and larger unit samples. **C)** Behavioral alignment of ANN models with macaques is uncorrelated with ImageNet top-1 accuracy across these models.

**Fig. S2:**
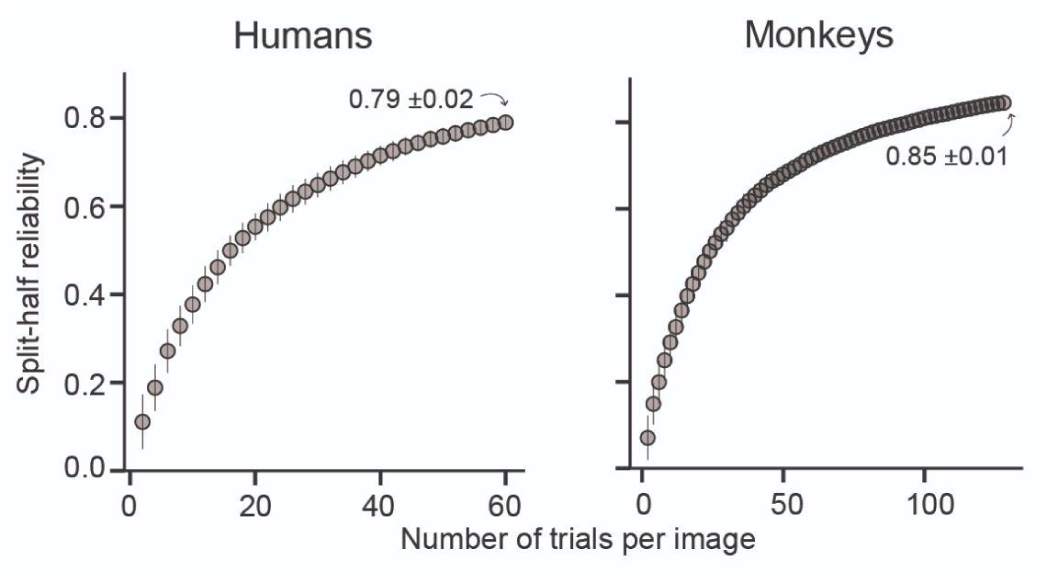
Split-half reliability of the image-level (B.I1) metric increases with repetitions per image. For humans (left) and monkeys (right) the split-half reliability was computed for an increasing number of trials. The error bars indicate the standard deviation across 100 random iterations.

**Fig. S3:**
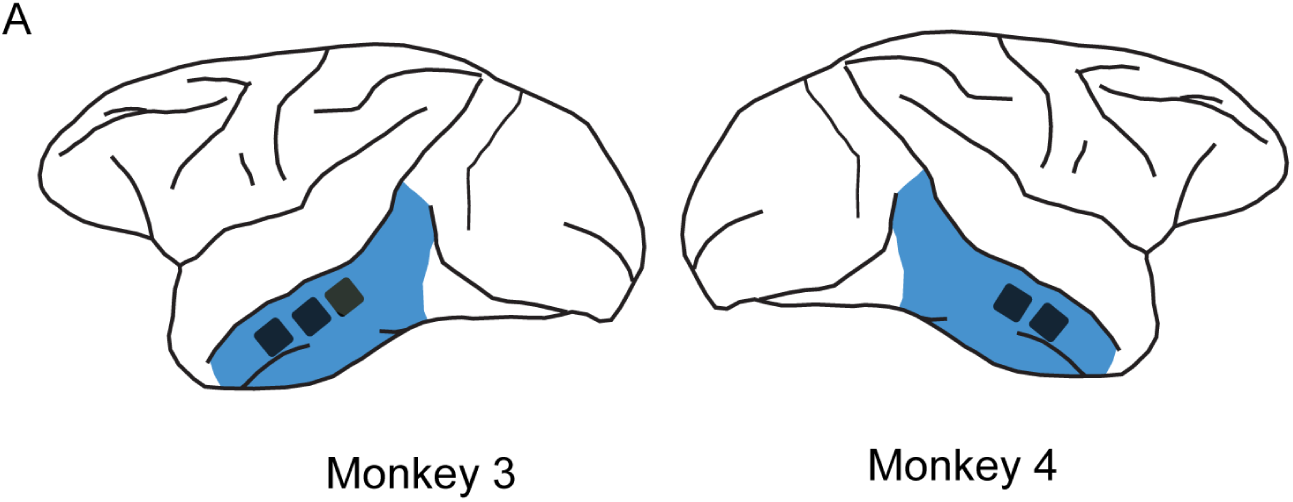
Location of implanted Utah array. Each black square marks an array and the blue area indicates an approximation of the portion of the IT cortex visible in this image.

